# TDP-43 deficiency links Amyotrophic Lateral Sclerosis with R-loop homeostasis and R loop-mediated DNA damage

**DOI:** 10.1101/2020.05.10.086652

**Authors:** Marta Gianini, Daisy Sproviero, Aleix Bayona-Feliu, Cristina Cereda, Andrés Aguilera

## Abstract

TDP-43 is a DNA and RNA binding protein involved in RNA processing and with structural resemblance to heterogeneous ribonucleoproteins (hnRNPs), whose depletion sensitizes neurons to double strand DNA breaks (DSBs). Amyotrophic Lateral Sclerosis (ALS) is a neurodegenerative disorder, in which 97% of patients are familial and sporadic cases associated with TDP-43 proteinopathies and conditions clearing TDP-43 from the nucleus, but we know little about the molecular basis of the disease. Here, we prove that mislocalization of mutated TDP-43 (A382T) in transfected neuronal SH-SY5Y and lymphoblastoid cell lines (LCLs) from an ALS patient cause R-loop accumulation, and R loop-dependent increased DSBs and Fanconi Anemia repair centers. Similar results were observed in a non-neuronal model of HeLa cells depleted of TDP-43. These results uncover a new role of TDP-43 in the control of co-transcriptional R-loops and the maintenance of genome integrity by preventing harmful R-loop accumulation. Our findings thus link TDP-43 pathology to increased R-loops and R loop-mediated DNA damage opening the possibility that R-loop modulation in TDP-43-defective cells might help develop ALS therapies.

## Introduction

TDP-43 is a nuclear RNA binding protein (RBP) with a repressor role of HIV-1 transcription. It binds to the trans-active response element DNA sequence of the viral genome [1,2]. Like other hnRNP proteins, TDP-43 binds to nascent pre-mRNA molecules when they are released from the RNA Polymerase II (RNApol II) and regulates RNA maturation either through sequential interactions with or in collaboration/antagonism with specific RNA binding factors [3]. TDP-43 is also involved in the regulation of non-coding RNAs like miRNAs and lncRNAs [4,5]. Thanks to its ability to recognize single-stranded DNA (ssDNA) or single-stranded RNA (ssRNA) with a preferential binding to (UG)n-enriched sequences [6], TDP-43 is involved in different steps of mRNA metabolism and in several mechanisms of genome integrity [7], consistent with the idea that RNA metabolism and DNA damage response (DDR) may be functionally interconnected [8].

Mutations in TDP-43 are associated to sporadic and familial cases of Amyotrophic Lateral Sclerosis (ALS), an adult onset, progressive neurodegenerative disease, caused by the selective loss of upper and lower motor neurons in the cerebral cortex, brainstem and spinal cord [9,10]. TARDBP is a major pathological gene for the ALS susceptibility and their mutations are found in 3% of familial and 2% of sporadic ALS cases [11,12]. Particularly, homozygous p.A382T TARDBP variation (A382T TDP-43) is one of the most common missense mutation in familial patients. A382T TDP-43 accumulation in the cytoplasm can reduce its physiological nuclear function, such as transcription regulation, mRNA splicing and transport [13,14,15] and miRNAs biogenesis [5,9]. Subsequent to this, the formation of oligomers and aggregates of TDP-43 in the cytoplasm may recruit native TDP-43 or other interactors proteins [16], constituting a gain of toxic function associated with neurodegeneration [17]. TDP-43 aggregates are identified as a major component of the ubiquitinated neuronal cytoplasmic inclusions deposited in spinal motor neurons both in familiar and sporadic ALS patients [18].

In addition to transcriptional autoregulation, TDP-43 can be cleaved into smaller C-terminal fragments before being enzymatically degraded to maintain its physiological levels [9,19] by a range of cysteine proteases, including caspases and calpains. Moreover, lines of evidence suggest that these CTFs can be produced via translation of an alternative transcript which is upregulated in ALS [20]. Recent studies proved that increased cytosolic sequestration of the poly-ubiquitinated and aggregated forms of mutant TDP-43 correlates with higher levels of DNA strand breaks, activation of DDR factors such as phospho-ataxiatelangiectasia mutated (ATM), phospho-53BP1, γH2AX in SH-SY5Y lines expressing wildtype (WT) or Q331K-mutant TDP-43 [21]. TDP-43 depletion leads to increased sensitivity to various forms of DNA damage and mutation in the C-terminus glycine-rich low-complexity region (LC domain) associates with the loss of its nuclear function [22]. In addition, TDP-43 colocalizes with active RNA polymerase II at sites of DNA damage along with the DDR protein, BRCA1, participating in the prevention and/or repair of R loop-associated DNA damage [23].

Evidence indicate that a major source of spontaneous DNA damage comes from the accumulation of R-loops, consisting in DNA-RNA hybrids and a displaced single strand DNA (ssDNA) [8]. Non-physiological R-loops occur as unscheduled events formed co-transcriptionally that can compromise genome integrity. Increasing evidence [16,24] has highlighted a common association of increased R-loops with a variety of genetic diseases, including neurodegenerative disorders, such as Amyotrophic Lateral Sclerosis (ALS) [25]. R-loop formation is enhanced in genomic regions containing highly repetitive DNA, which could facilitate the thermodynamic stabilization of RNA-DNA hybrids [26,27] and in cells mutated in genes encoding factors controlling R-loop homeostasis. Such factors are generally related to RNA processing and export or have DNA-RNA unwinding (helicase) or hybrid-specific ribonuclease (RNase H) activities [28,29]. However, a crucial role in prevention of R-loop formation is also played by the DDR. It is particularly notorious the role of BRCA2 and BRCA1 DSB repair factors or the Fanconi Anemia pathway (FA), especially FANCD2, involved in the repair of the inter-strand crosslinks (ICLs) and replication fork blockages [30, 31]. Deficiency on any of these factors lead to harmful R-loop accumulation in human cells [8].

All this, together with the fact that a number of neurodegenerative diseases highlight a particular sensitivity of the nervous system and motor neurons are associated with deficiencies in RNA metabolism and DDR, prompted us to investigate whether TDP-43 deficiency, as found in ALS cells, have a role in R-loop homeostasis that could explain previously described DDR defects of ALS cells. We show that TDP-43 plays a role in preventing R-loop accumulation and R loop-mediated DNA breaks in neuronal and non-neuronal cells and in patient cell lines, thus opening the possibility that R-loop modulation in TDP-43-defective cells might help develop ALS therapies.

## Results

### TDP-43 depletion leads to activation of DDR and of Fanconi Anemia pathway

A key regulatory role of TDP-43 in essential metabolic processes was previously suggested since silencing of TDP-43 in HeLa cells lead in dysmorphic nuclear shape, misregulation of the cell cycle, apoptosis, increase in cyclin-dependent kinase 6 (Cdk6) transcript and protein levels [40]. As a major readout associated with RNA transcription metabolic defects, we analysed accumulation of nuclear DNA-RNA hybrids in TDP-43 depleted HeLa cells (siTDP-43 HeLa cells).

Genomic DNA-RNA hybrids in siTDP-43 HeLa cells were first assessed by immunofluorescence microscopy (IF) using the anti-DNA-RNA hybrid S9.6 antibody, and determining the levels of the S9.6 signal in the nucleoplasm after subtracting the nucleolar contribution [41,42]. As controls we used HeLa cells transiently transfected with a mock control vector expressing GFP (siCTRL) or overexpressing the RNaseH1 enzyme, which specifically degrades the RNA moiety of hybrids [41]. A slight but significant increase of R-loops was observed in siTDP-43 HeLa cells in comparison to the siCTRL that was reduced upon RNaseH1 overexpression, which confirmed that S9.6 signal detected corresponded to R-loops (Fig 1A). Next, we determined R-loop accumulation by the more accurate method of DNA-RNA immunoprecipitation (DRIP)-qPCR, based specifically on the purification of genomic DNA-RNA hybrids of different sizes. In this case the S9.6 signal was determined for the highly expressed APOE and RPL13A genes, which have been established to be regions prone to form R-loops [31,41,43], and the SNRPN gene used as negative control [43,44]. We detected accumulation of DNA-RNA hybrids in the analysed genes in siTDP-43 HeLa cells compared to the siCTRL HeLa cells, obtaining a significative result on RPL13A gene. Importantly, RNaseH treatment induced a dramatically signal decrease confirming that it was R-loop dependent (Fig 1B).

**Fig 1.**
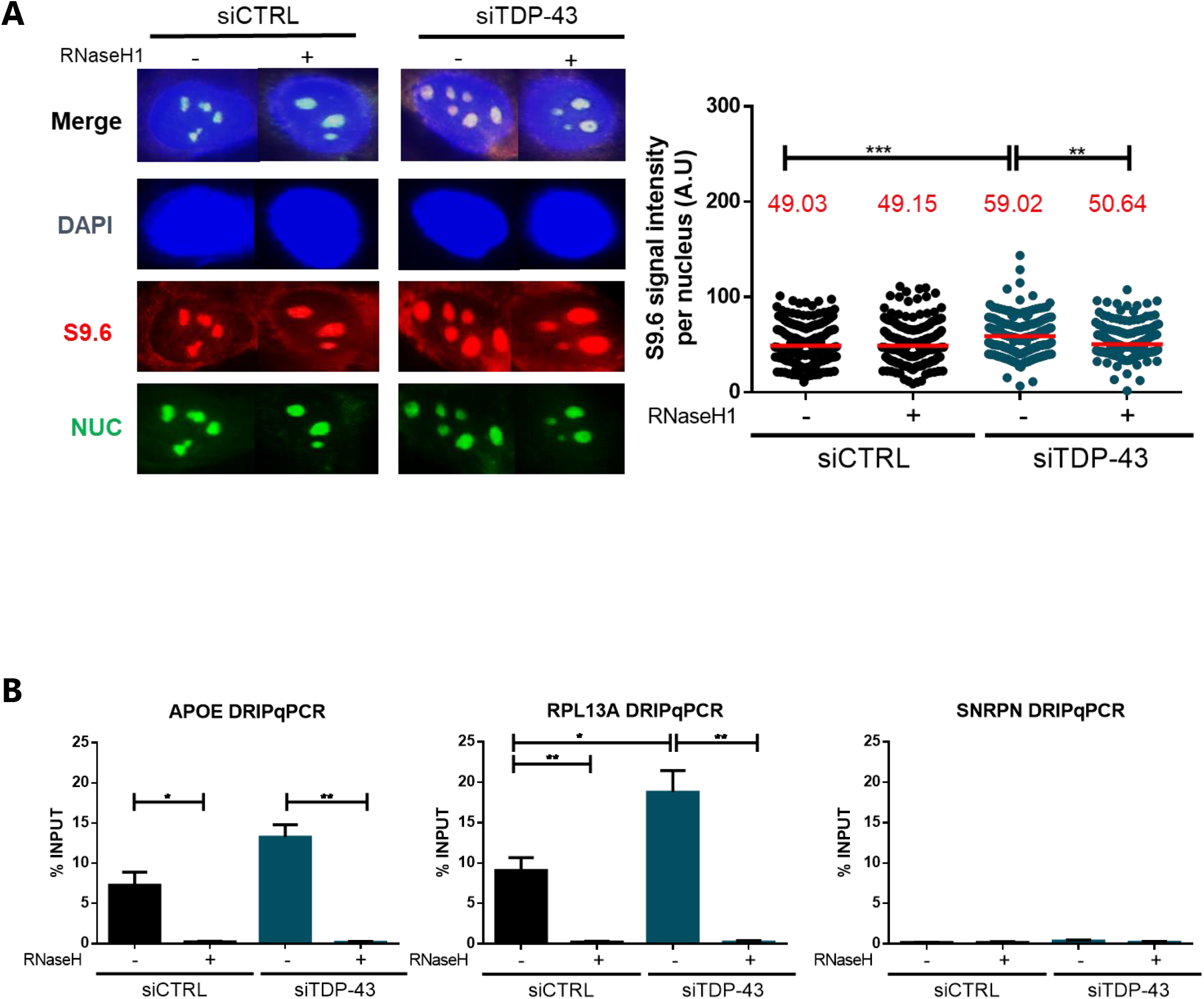
R-loops accumulation in siTDP-43 HeLa. **A)** siCTRL and siTDP-43 HeLa immunostaining with antiS9.6 antibody and anti-nucleolin antibody. The graph shows the median of the S9.6 intensity per nucleus after nucleolar signal removal. Around 300 cells from three independent experiments were considered. ***, P < 0,0002; **, P < 0,001 (Mann-Whitney U test, two-tailed). **B)** DRIP-qPCR using the anti S9.6 antibody at APOE, RPL13A and SNRPN genes are shown in siCTRL HeLa and siTDP-43 HeLa. Pre-immunoprecipitated samples were untreated (-) or treated (+) with RNaseH, as indicated. Data represent mean ± SEM from three independent experiments. *, P <0,04, **, P < 0,001 (Upaired t test, two-tailed).

Then, we investigated the functional impact of nuclear DNA-RNA hybrid enrichment on DDR, given that hybrids have been shown to enhance transcription-replication conflicts [45]. As can be seen in Fig 2A, γH2AX foci, as determined by IF, were significantly increased in siTDP-43 compared to siCTRL HeLa cells. γH2AX foci significantly decreased after RNaseH1 overexpression, indicating that the damage caused by TDP-43 depletion is R-loop mediated. It has been shown that the Fanconi Anemia (FA) repair pathway is a critical pathway to resolve R-loop mediated DNA breaks as the result of transcription-replication collisions and that the FA factors works at the collisions [30,31,34,46,47]. Therefore, we tested whether the damage generated by TDP-43 depletion was signalled by the FA pathway, for which we used the FANCD2 component [34]. As it can be seen in Fig 2B, FANCD2 foci were significantly increased in siTDP-43 HeLa cells compared to siCTRL HeLa. Importantly, this increase was reduced by RNaseH1 overexpression, proving that TDP-43 depletion is responsible for an activation of Fanconi Anemia repair factor caused by R-loop accumulation. The result is consistent with the idea that FANCD2 accumulates at R loop-containing sites at which the replication fork is blocked, similarly to inactivation of other RNA metabolic factors that lead to R-loop accumulation [32,34].

**Fig 2.**
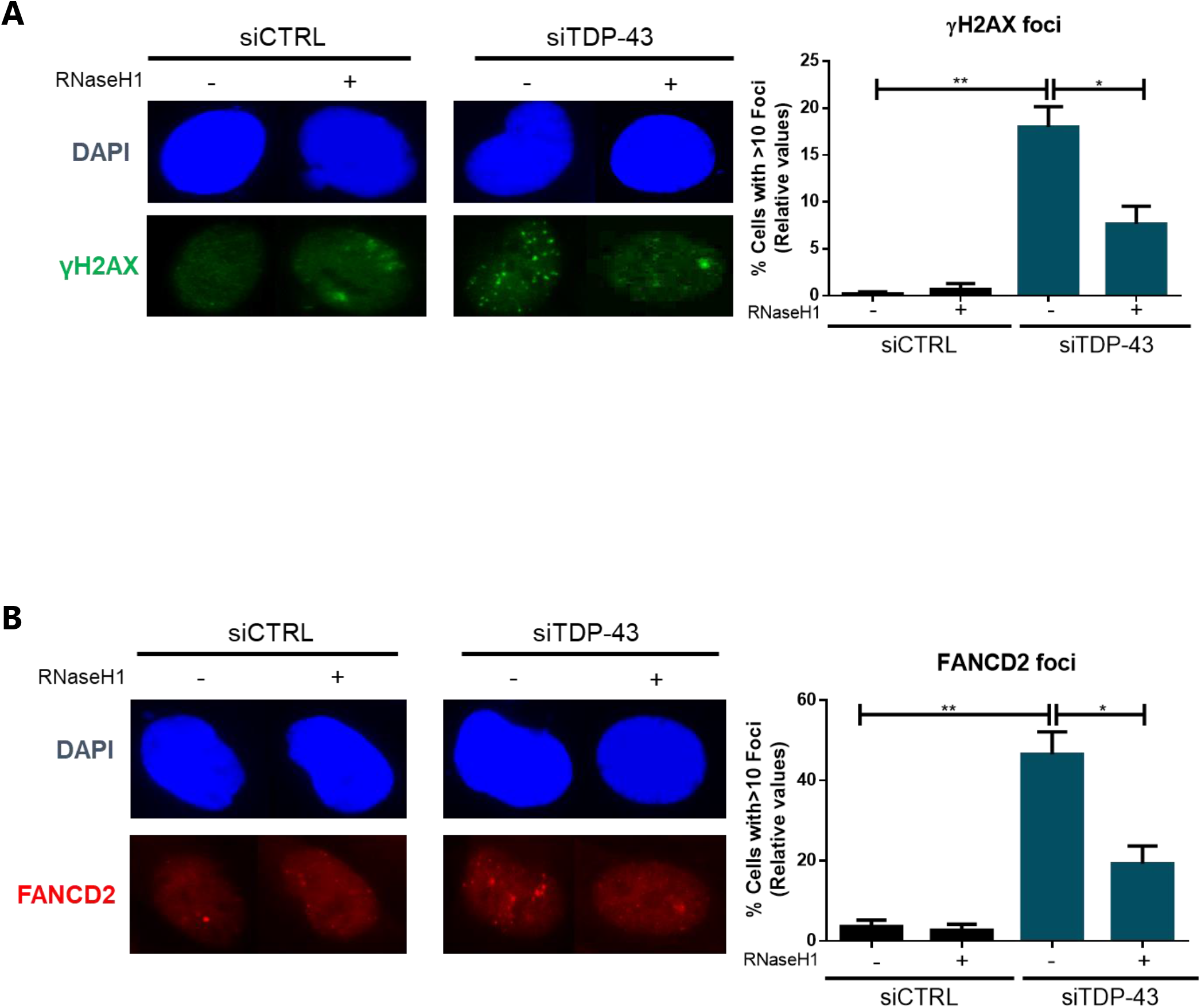
TDP-43 associates with FANCD2 and affects genome integrity in siTDP-43 HeLa. **A)** Detection of γH2AX foci by IF in siCTRL and siTDP-43 HeLa. The histogram shows the quantification of the relative amount of cells in percentage containing >10 γH2AX foci in each case. More than 100 cells were counted in each of the three experiments. Data represent mean ± SEM from three independent experiments. **, P<0,002 and *, P< 0,04 (Upaired t test, two-tailed). **B)** Detection of FANCD2 foci by IF in siCTRL and siTDP-43 HeLa. The histograms show the quantification of the relative amount of cells in percentage containing >10 FANCD2 foci in each case. Details as in **Fig 2A**. **, P<0,002 and *, P< 0,04 (Upaired t test, two-tailed).

### Cytoplasmic mislocalisation of mutated TDP-43 causes R-loop accumulation and leads to activation of DDR and of Fanconi Anemia pathway

In ALS patients harbouring TDP-43 mutations, TDP-43 mislocalises from the nucleus to the cytoplasm in detergent-resistant aggregated forms either full-length (43 KDa) and fragmented forms (35KDa, 25KDa), which can be ubiquitinated and hyperphosphorylated [48]. We hypothesized that TDP-43 mislocalisation due to missense mutations could have an impact on R-loop accumulation and DNA damage in ALS disease. To determine TDP-43 cellular localisation, we performed IF microscopy in basal SH-SY5Y, SH-TDP+ (overexpressing a GFP-tagged TDP-43 WT form) and SH-TDP382 (expressing the GFP-tagged p.A382T TDP-43 mutant form; SH) cells using an anti-TDP-43 or anti-GFP antibody, able to detect the wild-type nuclear protein and the cytoplasmic full length and fragmented forms. In SH-TDP+, the RBP was localised preferentially in the perinuclear area compared to non-transfected SH-SY5Y cells, in which localisation was predominantly nucleoplasmic. TDP-43 nuclear localization was significantly decreased in SH-TDP382 cells compared both to non-transfected SH-SY5Y and SH-TDP+, confirming that the A382T mutation is linked to TDP-43 cytoplasmic mislocalisation with formation of inclusions or aggregates (S1 Fig) as previously reported [49,50,51]. Results were the same when using the anti-TDP43 or anti-GFP antibodies.

Next, we tested whether mislocatisation of this RBP could impact onto genomic integrity and R-loop accumulation. We first assayed whether overexpression of wild-type TDP-43 and mutated and mislocalised TDP-43 affected R-loop accumulation. Since both mutated and overexpressed TDP-43 may affect its physiological role in miRNA biogenesis, we added in this case an ulterior treatment with RNaseIII, which degrades specifically dsRNAs, to counteract the reported ability of S9.6 to detect dsRNAs [35,52]. A significant increase of nucleolar S9.6 intensity was detected both in SH-TDP+ and SH-TDP382 cells compared to SH-SY5Y (Fig 3A). However, RNaseH1 overexpression caused nucleolar S9.6 signal decrease in SH-TDP382 cells only, but not in SH-TDP+ cells. The extra signal seen in TDP+ cells correspond to dsRNAs rather than DNA-RNA hybrids as shown by the sensitivity of such a signal to RNase III treatment. The reason why overexpression of TDP43 increases dsRNAs would need to be investigated further, and is not related to this study. Therefore, we conclude that mislocalisation of TDP-43 also causes R-loop accumulation.

**Fig 3.**
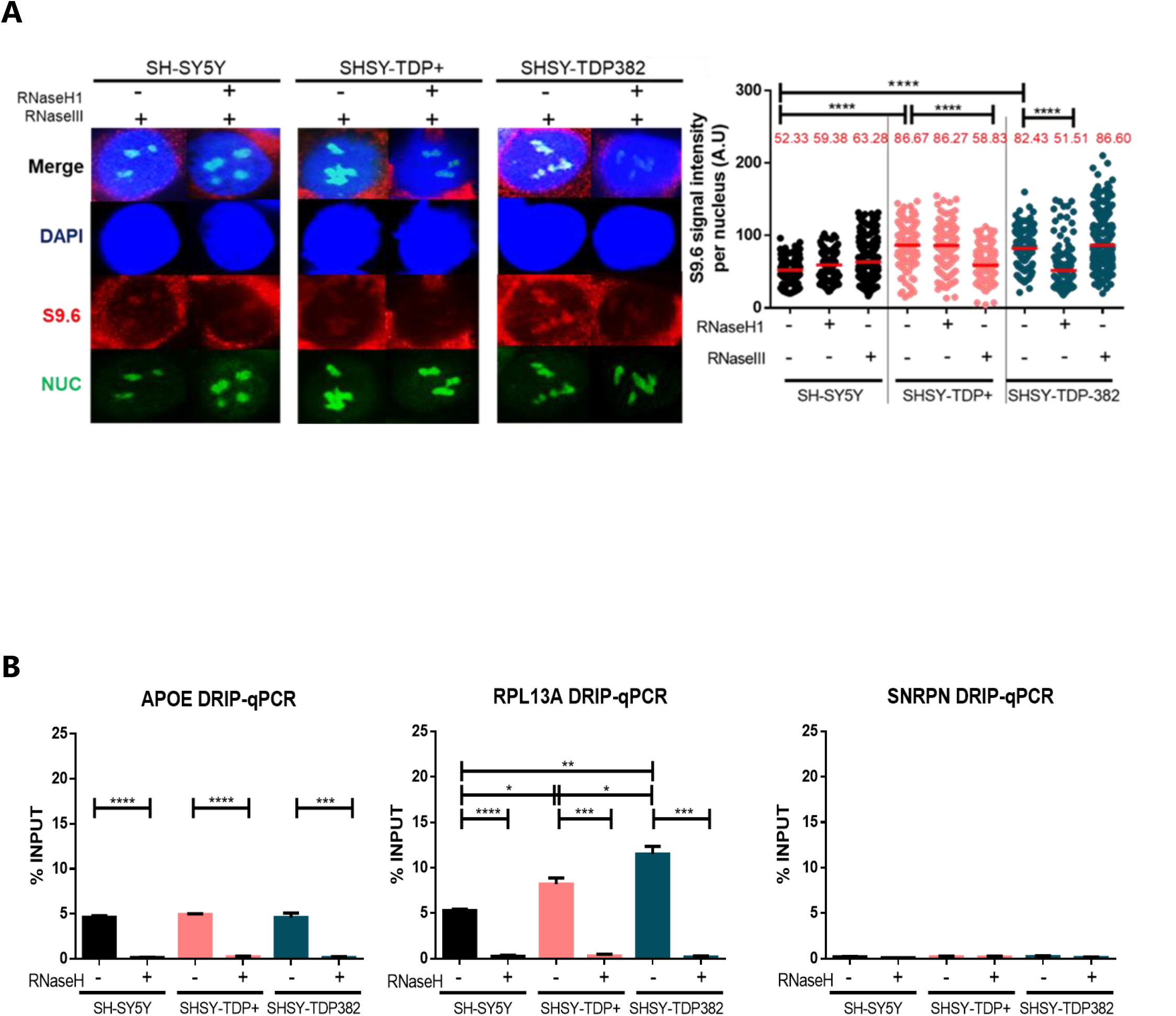
R-loops accumulation in SHSY-TDP382- and increased dsRNAs levels in SHSY-TDP+. **A)** IF using anti S9.6 antibody and anti-nucleolin antibody of SH-SY5Y, SH-TDP+ and SH-TDP382. The graph shows the quantification of S9.6 nucleolar signal. Details as in **Fig 1A**. ****, P < 0.0001 (Mann-Whitney U test, two-tailed). **B)** DRIP-qPCR using S9.6 antibody, at APOE, RPL13A and SNRPN genes in basal SH-SY5Y, SH-TDP+ and SH-TDP382. Details as in **Fig 1B**. *, P < 0,04; **, P < 0,002; *** P < 0,0003; ****, P < 0,0001 (Upaired t test, two-tailed).

We performed DRIP-qPCR in the neuroblastoma cell lines in the previously reported target genes [41,43,44], to confirm the results involving TDP-43 role in preventing R-loop accumulation in human cells. RNaseH treatment dramatically decreased the levels of the signal in all cases, confirming that the signal detected was specific for nuclear DNA-RNA hybrids. Whereas a significant R-loop accumulation was observed at the RPL13A gene in the SH-TDP382 mutant cells compared both to SH-SY5Y and SH-TDP+ (Fig 3B), this was not evident for APOE confirming the same tendency observed in siTDP-43 HeLa cells.

To test whether R-loop accumulation observed in SH-TDP382 cell lines causes DNA damage we determine the levels γH2AX foci by IF microscopy and their dependence of R-loops by testing whether RNaseH1 overexpression reduced them. As can be seen in Fig 4A, γH2AX foci were significantly increased in SH-TDP382 cells whereas this was not the case in SH-SY5Y and SH-TDP+ cells expressing the wild-type form of TDP-43. The increase in damage was suppressed by RNaseH1 overexpression, indicating that DNA break accumulation caused by mutant TDP43 was mediated by DNA-RNA hybrids.

**Fig 4.**
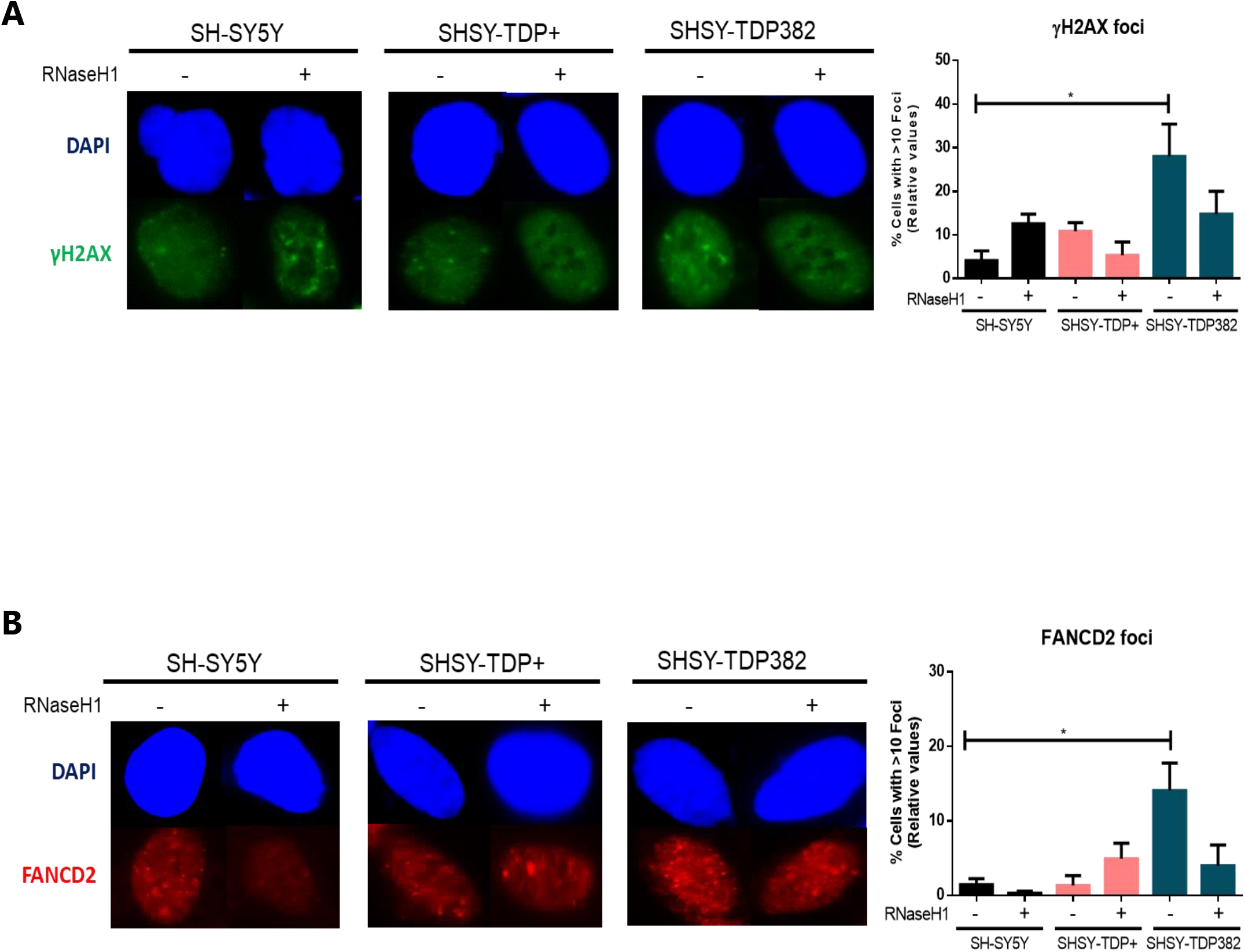
TDP-43 associates with FANCD2 and affects genome integrity in stable transfected SHSY-TDP382-. **A)** Detection of γH2AX foci, by IF in SH-SY5Y, SH-TDP+ and SH-TDP382. The histogram shows the quantification of the relative amount of cells in percentage containing >10 γH2AX foci in each case. Details as in **Fig 2A**. *, P < 0,04 (Upaired t test, two-tailed). **B)** Detection of FANCD2 foci by IF in SH-SY5Y, SH-TDP+ and SH-TDP382. The graph shows the quantification of the relative amount of cells in percentage containing >10 FANCD2 foci in each case. Details as in **Fig 2A**. *, P < 0,04 (Upaired t test, two-tailed).

Next, we tested whether the origin of such DNA damage was due to an increase in transcription-replication collisions enhanced by R-loops. We determined the levels of FANCD2 foci as previously reported. Notably, FANCD2 foci were significantly increased in SHSY-TDP382 mutant cells compared to SHSY-TDP+ and this increase was reduced by RNaseH1 overexpression (Fig 4B). Therefore, the ALS pathogenic TDP-43 mutation in the analysed neuronal model leads to a comparable functional effect to that observed in silenced HeLa cells. The pathogenic TDP43 mutation causes an increase in DNA breaks derived from R-loop accumulation that promotes transcription-replication collisions that are processed by the FA pathway, as reported for other cases of recombinogenic R-loops [30,31,34,46,47].

### Accumulation of R-loops in p.A382T TDP-43 mutated lymphoblastoid cell lines

Next, we used lymphoblastoid cell lines (LCLs) from a TDP-43 mutated patient carrying p.A382T mutation (LCL-TDP382), a sporadic ALS patient (LCL-SALS) and an healthy control (LCL-CTL) to confirm the role of TDP-43 in R-loops removal in ALS. We performed IF microscopy for S9.6 and TDP-43 in the three cell lines mentioned with two fixation methods, methanol (S2 Fig) and paraformaldehyde (Fig 5A). R-loop quantification of the dot blot confirmed a significant R-loop accumulation in LCL-TDP382 cells using both fixation methods. Paraformaldehyde fixation guaranteed a decreased detection level of TDP-43 signal in the nucleus of LCL-TDP382 in comparison to LCL-CTL and LCL-SALS, which is in accordance with the mislocalisation of the mutated protein in the cellular cytoplasm, causing the loss of its nuclear physiological function [53]. Moreover, there is a colocalization of S9.6 signal with TDP-43 in the perinuclear area of LCL-TDP382 cells in comparison to LCL-CTL and LCL-SALS. R-loop quantification was also determined in LCLs by flow cytometry, in which case the analysis reported an increased positivity of S9.6 intensity in the orange peak associated with LCL-TDP382, in comparison to the blue peak associated with LCL-CTL (Fig 5B). The positive signal in LCL-TDP382 represented by the orange peak was clearly suppressed by RNase H treatment in the same sample detected as green peak, confirming that the detected signal corresponds to DNA-RNA hybrids (Fig 5C). S9.6 mean fluorescence of LCL-TDP382 in a triplicate experiment was significantly increased compared to LCL-CTL and LCL-SALS, and this increase was reverted by RNaseH treatment (Fig 5D).

**Fig 5.**
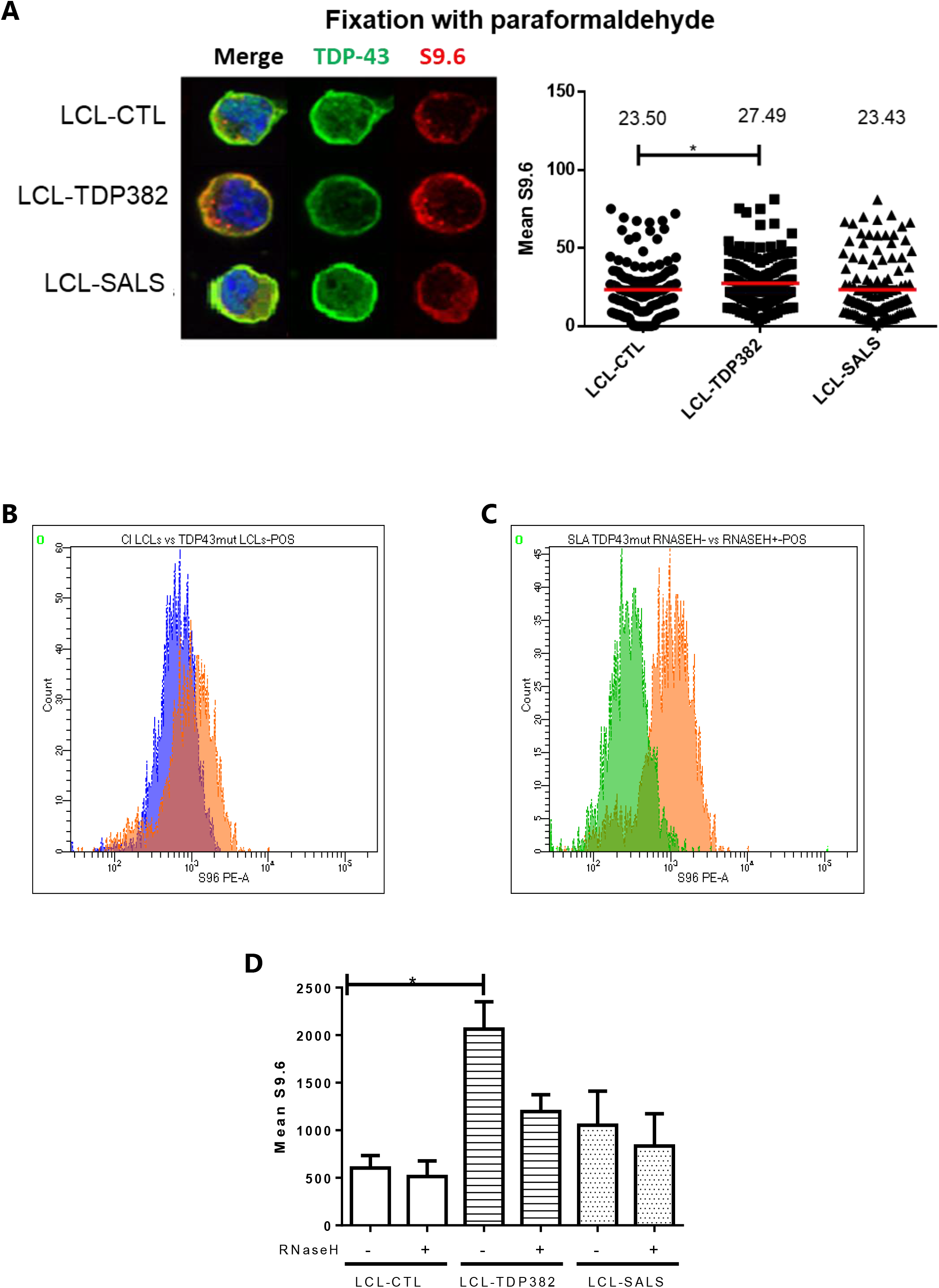
R-loops accumulation in p.A382T TDP-43 mutated LCLs. **A)** IF of LCL-CTL, LCL-TDP382, LCL-SALS using an anti-TDP-43 antibody and an anti-S9.6 antibody after paraformaldehyde fixation. **B)** The dot blots show an increase of S9.6 signal intensity *, P< 0,0385 (Mann-Whitney U test, two-tailed). **C)** Flow cytometry plot reports the amount of R-loops in LCLs (blue) in comparison to LCL-TDP382 (orange). **D)** The RNaseH action on LCL-TDP382 decreases the presence of R-loops signal (orange peak vs green peak). **E)** Histogram of S9.6 mean fluorescence of LCL-CTL, LCL-TDP382, LCL-SALS in presence (+) and in absence (-) of RNaseH proved significative R-loops accumulation in LCL-TDP382. The value represented is the mean ± SEM of three biological experiments. ANOVA, Newman-Keuls Multiple Comparison Test, *P <0,05.

Finally, we investigated the possibility that TDP43 could have a role on DNA-RNA hybrids directly, in which case we should expect some kind of physical association. Therefore, we wondered whether TDP-43 and genomic DNA-RNA hybrids colocalize by performing a co-immunoprecipitation (coIP) in chromatin (Chr) fractions from the three cell lines. At the same time, we extracted whole lysate (WL) fractions from the same samples as control for the cytoplasmatic fraction (Fig 6A-B). In the Chr fraction, coimmunoprecipitation could be observed with the S9.6 antibody. In LCL-TDP382, the TDP-43 mutant protein showed lower levels of co-IP, while in the WL fraction of the same sample the co-IP signal was similar to WT, which suggests that the mutant full length TDP-43 was not able to interact with R-loops due to its sequestration at the cytosolic compartment in the cell [53], as previously seen in Fig 5A. Interestingly, the truncated TDP-35 form detected in Chr fraction of LCLs show high levels of S9.6 co-IP in the WL fraction of LCL-TDP382 in comparison with control LCL-CTL and LCL-SALS. This specific C-terminal mutation may predispose TDP-43 to fragmentation into CTFs, which as reported in literature are transported out of the nucleus and accumulated into complexes with RNA transcripts [49]. The result suggests that a fraction of cellular TDP-43 is present in chromatin in association with DNA-RNA hybrids and both wild-type and truncated forms can be found in the cytoplasm, Therefore, we can conclude that TDP-43 has a role in RNA metabolism from nucleus to cytoplasm that help prevent cells to accumulate co-transcriptional harmful R-loops.

**Fig 6.**
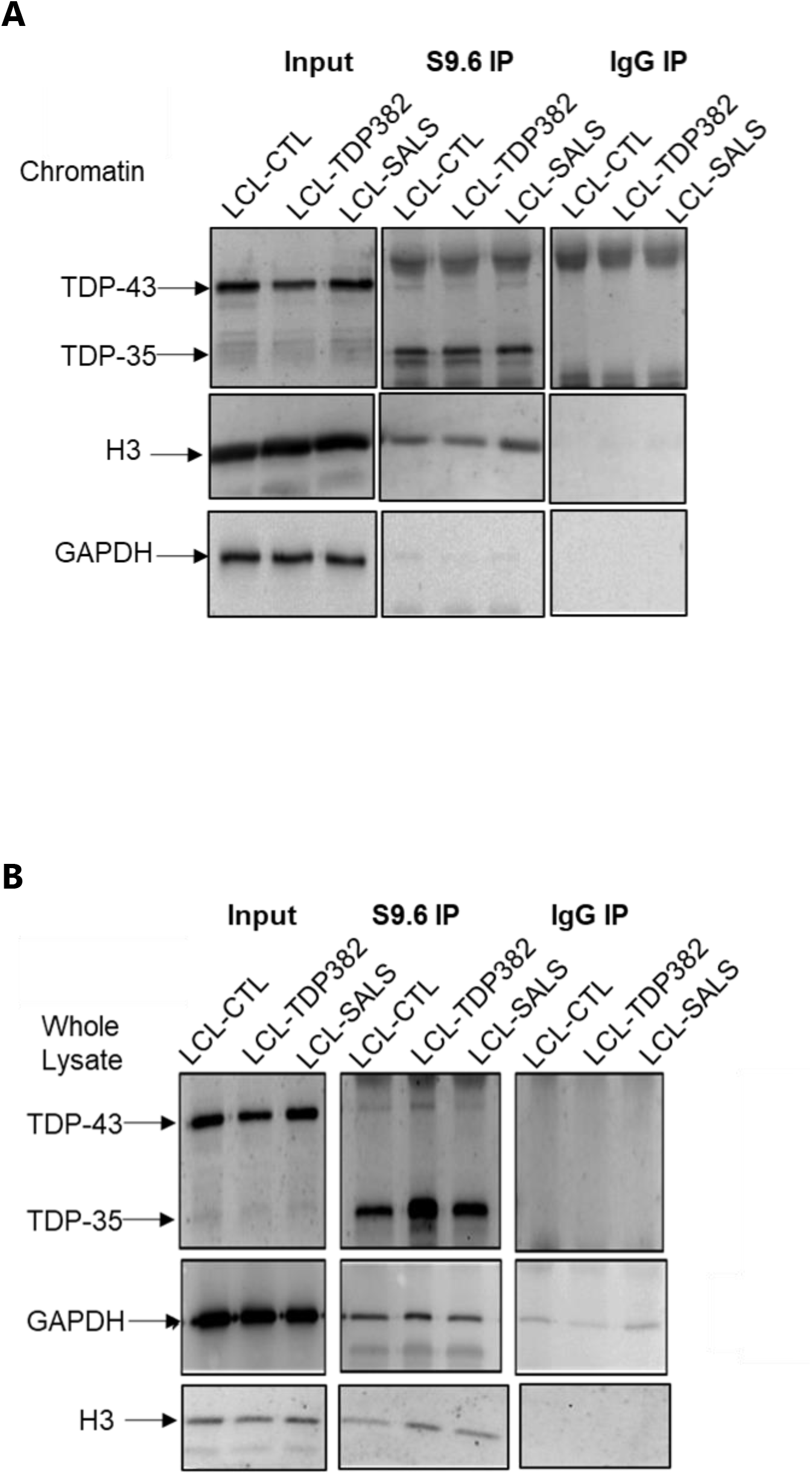
TDP-43 and TDP-35 strongly interact with S9.6 antibody in TDP-43 mut LCLs WL fraction. **A and B)** CoIP between S9.6 and TDP-43 in chromatin of LCL-CTL, LCL-TDP382, LCL-SALS. Input, S9.6 IP and IgG IP of chromatin fraction were loaded on a 10% SDS-PAGE and then immunoblotted with TDP-43, H3 and GAPDH as nuclear and cytosolic loading control. S9.6 binding was tested by qPCR.

## Discussion

Mislocalisation of TDP-43 causes a gain of neurotoxic function characteristic of the neurodegeneration process in ALS patients [54]. Moreover, TDP-43 aggregation induced by double-site mutations and TDP-43 knockdown have a common set of differentially expressed proteins, behaving in a similar way [55]. Here we show that mislocalisation of mutated TDP-43 can sequester full length TDP-43 form in cytoplasmic inclusions preventing its physiological nuclear function. Importantly, this function is related to R-loop homeostasis. Nuclear depletion of TDP43, as achieved by either mislocalisation to the cytoplasm or siRNA depletion in different cell types, causes a significant increase in harmful R-loops that leads to DNA breaks and FANCD2 foci. The results suggest that the TDP-43 RNA-binding protein has a key role in preventing R-loop accumulation as a safeguard of genome integrity.

Silencing TDP-43 by siRNA in HeLa cells led to a significative increase of R-loop signal by S9.6 IF compared to the siCTRL control. This was confirmed by reversion of signal in case of RNaseH1 overexpression. DRIP-qPCR revealed an important R-loop presence on the tested RPL13A gene encoding a highly expressed ribosomal protein involved in pathways of viral mRNA translation and of rRNA processing, whose impairment could lead to alteration of protein homeostasis as well as modifications of RNA metabolism [56]. It is worth noting that TDP-43 interacting proteins largely cluster into two distinct interaction networks and RPL13A in particular is implicated in TDP-43 cytoplasmic interactome cluster, resulting fundamental for RNA translation regulation [57]. Also, experimental studies reported that both C9orf72 mut ALS patients’ derived iPSCs and TDP43-EGFP overexpressing iPSCs presented a set of commonly destabilized RNAs involved in the ribosomal and in the oxidative phosphorylation pathways. As consequence of RNA instability in iPSCs it was detected an increase of cytoplasmic ribosome proteins including RPL13A, suggested as cellular compensatory response for preservation of protein synthesis capacity. Moreover, the analysis of 3’UTRs of transcripts in the same cells showed a high enrichment in motifs recognized by RBPs and involved in the formation of cytoplasmic inclusions, that in turn exhibited an alteration of their stability [58]. We do not believe that the particular effect observed in accumulation of R-loops in RPL13A is related to these phenotypes, since ribosomal protein genes are highly transcribed, thus favouring R-loop formation. However, it is certainly possible that indirectly R-loop accumulation itself in ribosomal protein genes, may be linked to the misbehaviour of the translation and UTR function.

The absence of nuclear TDP-43 affects the DDR, consistent with previous reports [23]. We showed that siTDP-43 HeLa cells had a significative increase of DSBs as determined by γH2AX foci. Importantly, this DSB increase was R-loop-dependent, as it could be fully reverted by RNaseH1 overexpression, and it was accompanied by an accumulation of FANCD2 foci that was also R loop-dependent. The result indicates that TDP-43 prevents the co-transcriptional accumulation of harmful R-loops that promote transcription-replication conflicts that have to be resolved by the FA pathway, consistent with the previously reported role for the FA pathway [45].

Notably, our assay of the impact of the TDP-43 pathogenic ALS mutation and overexpression in a neuroblastoma cell line, SH-SY5Y, revealed that the pathogenic A382T mutation in SH-SY5Y cells also affected the TDP-43 role controlling R-loops homeostasis, leading to a higher detection of genomic RNA-DNA hybrids, as detected by IF and DRIP-qPCR. As expected DSBs and replication blockage detected by γH2AX and FANCD2 foci, respectively, were also increased in an R loop-dependent manner.

The effect of TDP-43 deficiency in R-loop homeostasis could be related to the accumulation of aberrant transcripts and hybrids that are trapped in persistent RNA-processing foci present in the cytoplasmic compartment of cell previously reported [59]. Our analysis of ALS patient-derived LCLs, a valid cellular model to understand ALS disease [51], also revealed an increased level of nuclear RNA-DNA hybrids in LCL-TDP382 as well as co-localization of S9.6 antibody with a fraction of TDP-43 in the perinuclear area. It is possible that either sequestering of the misfolded and mutated form of TDP-43 in inclusions could cause a loss of its nuclear function or the formation of TDP-43 aggregates in the cytoplasm could recruit native TDP-43 or other interactors proteins constituting a gain of toxic function [59]. The detection of higher S9.6 signal in both in LCL-SALS and of LCL-TDP382 underlines that R-loops may be a general condition in ALS, potentially improving RNA metabolism dysregulation and neurotoxicity that appear to be major contributors to the pathogenesis of this neurodegenerative disease.

It is worth noticing that when TDP-43 was overexpressed (SH-TDP+), a significant increase of an S9.6 signal that was sensitive to RNase III treatment but not to RNaseH1 overexpression, in contrast to A382T mutant. Knowing that S9.6 can also detect dsRNAs [35,52] this result indicates that TDP-43 overexpression leads to an accumulation of dsRNA molecules. Interestingly, TDP-43 has been reported to co-localize with Dicer and Ago2, but their interaction is inhibited by aggregates formation in response to cellular stressors or also by overexpression of human TDP-43 [60]. There is no evidence that overexpression and mutation of TDP-43 lead to double hairpin pre-miRNA accumulation in neuronal models, so it is formally possible that dsRNA accumulation observed in SH-SY5Y overexpressing TDP-43 could be associated to inhibition of Dicer processing function responsible for the loss of maturation of pre-miRNAs, presented as dsRNAs hairpin structure, in mature miRNAs. However, it may also be possible that excess of RBPs would prevent normal RNA metabolism by excess of cellular RBPs that would bind to any RNA molecule having secondary structure segments. A negative impact of RBP overexpression on RNA metabolism has been reported in other cases [37]. These are possibilities to explore in the future.

The chromatin fraction of TDP382 mutated LCLs showed a weaker association between TDP-43 and the S9.6 signal not observed in whole cell extracts, suggesting a lower TDP-43 presence in the nucleus and/or lower capacity of this mutant form to associate with R-loops. However, the S9.6 signal detected in the chromatin fraction of LCLs by CoIP suggested an important role of the TDP-35 truncated form. It is known that in ALS pathological conditions TDP-43 can generate CTFs such as 35 kDa fragments both upon cleavage by caspases at intrinsic caspase cleavage sites [11], both by via translation of upregulated alternative transcript [20]. Due to the lack of nuclear localization signal (NLS), CTFs TDP-35 mislocalise to the cytoplasm, where may associate with RNA forming cytoplasmic inclusions [61]. Indeed, the biochemical analysis suggests that TDP-35 facilitates aggregate assembly promoting inclusion formation [62] and might transport different types of RNA structures. It is known that TDP-35 can also recruit full-length TDP-43 to cytoplasmic deposition from functionally nuclear localization [63] and TDP-43 continuously shuttles between nucleus and cytoplasm in a transcription-dependent manner [64]. The higher S9.6 signal in the whole lysate fraction of LCL-TDP382 compared to the other LCLs likely may reflect inclusions formed by dsRNAs.

It is becoming clear that impaired RNA regulation and processing is a central feature in ALS pathogenesis. Our study, reinforces the need of understanding the specific role of ALS in RNA metabolism, and in particular in cells defective in the TDP-43 RBP, beyond its effect on the formation of RNA inclusion bodies in the cytoplasm. Even though this is a common read-out of ALS, our study showing an increase in genomic R-loops and DNA damage and Fanconi Anemia foci, expected for obstacles blocking replication, suggests that an important cause of the disease may be linked to impairment of nuclear RNA biogenesis and impact on DDR. The application of RNA-based therapies to modulation of gene and subsequent protein expression is an attractive therapeutic strategy, that could be considered in the future for the treatment of ALS and other neurodegenerative diseases.

## Materials and methods

### Human cell culture and transfection

HeLa cells (CCL-2, American Type Culture Collection [ATCC]) and SH-SY5Y (CRL-2266, American Type Culture Collection [ATCC]) were cultured in Dulbecco’s modified Eagle medium (DMEM, Thermo Fisher Scientific) supplemented with 10% fetal bovine serum (FBS) (S1710-500, Biowest), 1% Penicillin/Streptomycin (L0022-100, Biowest), and 1% L-glutamine (11539876, Gibco). Murine hybridoma cell line ATCC^®^ HB-8730™ were grown in DMEM supplemented with 10% fetal bovine serum, 4 mM L-glutamine, 4500 mg/L glucose, 1 mM sodium pyruvate, and 1500 mg/L sodium bicarbonate (5% CO^2^). Lymphoblastoid cell lines (LCLs) were harvested with RPMI 1640 (42401018, Gibco) supplemented with 20% of FBS, 1% Penicillin/Streptomycin and 1% L-glutamine.

HeLa cells were transiently transfected using DharmaFect (Dharmacon) according to the manufacturer’s instructions with siCTRL (D-001810-01-05) and siTDP-43 (L-012394-00-0005) and collected after 72h, as described by Dominguez-Sanchez et al. [32]. HeLa cells and SH-SY5Y were stably transfected using Lipofectamine 2000 (Invitrogen, Carlsbad, CA), according to the manufacturer’s instructions. HeLa cells were transfected with the following plasmids: pEGFP as control vector expressing GFP (Clontech), and pEGFP-M27, containing the GFP-fused RNaseH1 lacking the first 26 amino acids responsible for its mitochondrial localization cloned into pEGFP [33]. All assays were performed 72 h after siRNA transfection or 24 h after plasmid transfection [32].

SH-SY5Y cells were stably transfected using Fugene HD Transfection Reagent (Promega) with C-terminally GFP-tagged TDP-43 WT vector (PS100010 pCMV6-AC-GFP) and C-terminally GFP-tagged TDP-43 A382T mutation vector (CW303334 Mutate ORF of RC210639 at nt position 1144, to nr change G to A,to achieve A382T, keep insert in PS100001), purchased from OriGene. During the paper the SH-SY5Y are respectively named: SH-SY5Y expressing basal levels of TDP-43, SH-TDP+ overexpressing a GFP-tagged TDP-43 WT form and SH-TDP382 expressing the GFP-tagged p.A382T TDP-43 mutant form.

### Immunofluorescence microscopy

For S9.6 IF analysis in HeLa and SH-SY5Y, cells were fixed with cold methanol for 10 minutes at −20°C according the literature [34]. SH-SY5Y cells were treated with 40 U/ml RNaseIII (1 U/μl, Thermo Fisher Scientific) for 30 minutes at 37° using 1X RNase III Reaction Buffer. For γH2AX and FANCD2 IF analysis in HeLa and SH-SY5Y, cells were incubated with a fixation solution (PFA 4%, Triton-X 0,1%) for 10 minutes at room temperature (RT) as previously described [35]. For S9.6 and TDP-43 IF microscopy analysis in LCLs, cells were fixed with two methods: one using formaldehyde and cold acetone and the other using cold methanol.

The following antibodies were used: anti-nucleolin antibody (ab50279, Abcam), S9.6 monoclonal antibody (ATCC^®^ HB-8730™ hybridoma),anti-TDP-43 antibody (Clone: 6H6E12, Proteintech), anti-γ-H2AX antibody (ab2893, Abcam), anti-FANCD2 antibody (sc-20022, SantaCruz), anti-H3S10P antibody (06-570, Sigma-Aldrich). Antibody signals were detected on a Leica DM6000 microscope equipped with a DFC390 camera (Leica). Data acquisition was performed with LAS AF (Leica). R-loops signal in the nucleoplasm of HeLa and SH-SY5Y cell lines was quantified using ImageJ program by measuring the S9.6 integrated density observed in the DAPI-stained nucleus, subtracting the nucleolar contribution, detected by nucleolin antibody. γH2AX and FANCD2 foci were determined in IF of HeLa and SH-SY5Y cell lines through the relative number of cells containing >10 foci in the nuclei in each condition. In each IF experiment we maintained around 100 counted cells for improving their comparability.

### DNA-RNA immunoprecipitation-qPCR

DNA-RNA immunoprecipitation (DRIP) was performed on HeLa and SH-SY5Y cells as already described in literature [36,37]. The amount of R-loop levels was quantified as a function of input DNA, that for each sample was 10% of the entire amount.

### Flow Cytometry analysis

LCL control (LCL-CTL), LCL sporadic patient (LCL-SALS) and LCL A382T TDP-43 mutated patients (LCL-TDP382) were incubated with a vitality dye for 15 minutes (Zombie Violet™ Fixable Viability Kit, BioLegend). Then, LCLs were incubated for 20 minutes with anti-CD19 antibody for B lymphocytes recognition. Cells were fixed and permeabilized using a kit based on saponin permeabilization (Fixation/Permeabilization Solution Kit, BD) following a protocol described by Schauer U. et al. [38]. As negative control for R-loop presence, cells were treated with 60 U/ml of ribonuclease H (RNase H, 5.000 units/ml, NEB) using RNase H buffer at 37°C for 1 hour. As negative control for ssRNAs, cells were treated with 100 μg/mL ribonuclease A (RNase A, 10 mg/mL, Thermofisher) in 0.3M NaCl buffer at 37°C for 1 hour. At the end, cells were stained for one hour with conjugated anti-S9.6 antibody (PE/R-Phycoerythrin Conjugation Kit, Abcam) and analysed by flow cytometry (BD FACS Canto II). Logarithmic amplification was used for all channels and FACSDIVA was used for the analysis.

### Co-immunoprecipitation studies

Chromatin fraction (Chr fraction) and whole lysate fraction (WL fraction) were extracted from LCL-CTL, LCL-SALS and LCL-TDP382 cells. Cells (10^7^) were resuspended in buffer A (25 mM HEPES, 1.5 mM MgCl2, 10mM KCl, 0.5% NP-40, 1 mM DTT) incubated on ice for 10 minutes to isolate the nuclei and spin at 5000 rpm for 5 min at 4 °C. The pellet was resuspended in buffer B (50 μM HEPES pH 7.8, 140 mM NaCl2, 1mM EDTA, 1%Triton X-100, 0.1% Na-deoxycholate, 0.1% SDS) and sonicated for 5 cycles (30 ON/30 OFF) using a Bioruptor ^®^ Sonicator Brochure – Diagenode. All buffer were supplemented with protease and phosphatase inhibitor. Magnetic beads (Dynabeads, Invitrogen) were conjugated with 10 μg S9.6 antibody and respective murine IgG antibody (sc-2025, Santa Cruz) for 2h at 4°C in agitation, followed by incubation O/N with the samples, recovering 10% prior to incubation. After washes, a part of the samples was eluted using elution buffer (50mM Tris pH 8.0, 1mM EDTA, 1% SDS) at 65°C for 10 minutes twice and resuspended in TE buffer for qPCR analysis. S9.6 immunoprecipitation was evaluated by qPCR in genomic sites enriched for R-loop presence in these cells (NOP58 and ING3) [39]. Input (10% fraction for each sample), S9.6 and IgG were loaded on a 10% SDS-PAGE and co-immunoprecipitation was evaluated by immunoblotting with rabbit anti-TDP-43 (1:1000 dilution), rabbit anti-H3 (1:1000 dilution), rabbit anti-GAPDH (1:10000 dilution). Anti-TDP-43 antibody (Clone: 6H6E12, Proteintech Europe), anti-H3 antibody (ab1791, Abcam), anti-GAPDH antibody (GTX100118, GeneTex), IgG antibody (sc-2025, Santa Cruz) were used.

### Statistical Analysis

Statistical analysis was performed by Student t-test and by One-Way Analysis of Variance (ANOVA test) followed by post hoc comparison as a post-test (GraphPad Prism version 5, San Diego, CA, USA), unless otherwise specified. Values were considered statistically significant when p values were < 0.05.

## Acknowledgements

Research in C.C’s lab was funded by Fondazione Regionale per la Ricerca Biomedica for TRANS–ALS (Translating Molecular Mechanisms into ALS risk and patient’s well-being: FRRB 2015-0023) and in A.A.’s lab was funded by grants from the European Research Council (ERC2014-AdG669898 TARLOOP), the Spanish Ministry of Economy and Competitiveness (BFU2016-75058-P), and the European Union (FEDER). A.B was supported by a Juan de la Cierva-Formación fellowship (FJCI-2017-34536) from Spanish Ministry of Science and Innovation. M.G. was recipient of a scholarship provided by Erasmus+ Mobility for Traineeships from University of Pavia.

## Authors’ contributions

The work was designed by C.C. and A.A and supervised by D.S. and A.B-F. All experiments were performed by M.G. and data analysed by M.G., D.S. and A.B.-F The paper was written by M.G., D.S, C.C and A.A. All authors discussed, revised and agreed with the final version.

## Supplemental Figures

**S1 Fig. Cytoplasmic mislocalization of TDP-43 in p.A382T TDP-43 SH-SY5Y.** TDP-43 immunostaining in not transfected SH-SY5Y, SHSY-TDP+ and SHSY-TDP382-. Enlargement of TDP-43 WT SH-SY5Y underlined TDP-43 perinuclear localization, while white arrows TDP-43 A382T SH-SY5Y made more evident cytoplasmic TDP-43 inclusions. The dot blot shows the median of the TDP-43 signal intensity per nucleus. Around 300 cells from two independent experiments were considered. ****, P < 0.0001 (Mann-Whitney U test, two-tailed).

**S2 Fig. R-loops accumulation in p.A382T TDP-43 mutated LCLs.** IF of LCL-CTL, LCL-TDP382, LCL-SALS using an anti-TDP-43 antibody and an anti-S9.6 antibody after methanol fixation. The dot blots show an increase of S9.6 signal intensity *, P< 0,0385 (Mann-Whitney U test, two-tailed).

