## Supplementary figures and images for "TDP-43 deficiency links Amyotrophic Lateral Sclerosis with R-loop homeostasis and R loop-mediated DNA damage"

### Supplemental Fig. 1

Supplemental Figure 1. Giannini et al.

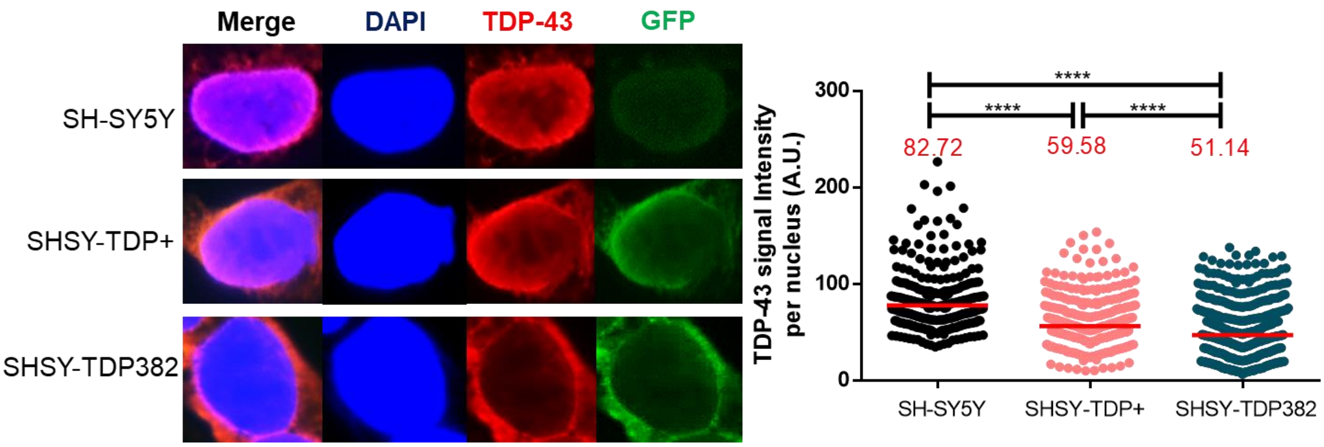

### Supplemental Fig. 2

Supplemental Figure 2. Giannini et al.

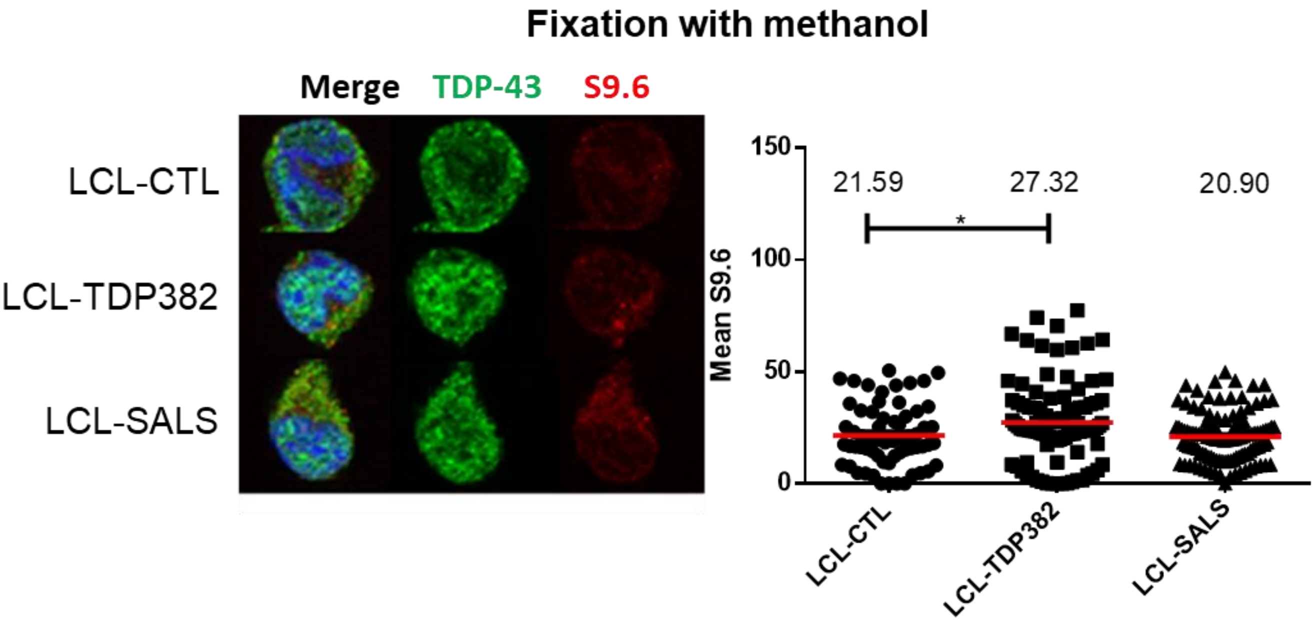
